# Reaching reproduction in a large carnivore: role of early environment and intrinsic traits

**DOI:** 10.1101/2025.01.24.634662

**Authors:** Léa Auclair, Cecilia Di Bernardi, Håkan Sand, Mikael Åkesson, Barbara Zimmermann, Øystein Flagstad, Petter Wabakken, Camilla Wikenros

**Affiliations:** National Museum of Natural History, Centre d’Écologie et des Sciences de la Conservation (CESCO), 57 Rue Cuvier, 75005 Paris, France; Grimsö Wildlife Research Station, Department of Ecology, Swedish University of Agricultural Sciences, SE-739 93 Riddarhyttan, Sweden; Faculty of Applied Ecology, Agricultural Sciences and Biotechnology, Inland Norway University of Applied Sciences, Evenstad, NO-2418, Elverum, Norway; Norwegian Institute for Nature Research (NINA), P.O. Box 5685, Torgarden, NO-7485, Trondheim, Norway

**Keywords:** Canis lupus, juvenile stage, life-history traits, natal territory, reproduction

## Abstract

To reach reproduction, individuals must survive the juvenile stage, a critical period of low survival rates in large carnivores. We analysed data from 582 wolves (*Canis lupus*) identified by DNA during their first year in Sweden and Norway, to investigate intrinsic and extrinsic factors within the natal territory affecting the probability to reach reproduction, i.e. having pups surviving at least 5 months of age. Factors included main prey density, road density, human density, and proximity to non-breeding zones, as well as sex, inbreeding and being collared. Of the 582 wolves identified, 21% reached reproduction. Human density and whether a wolf was collared were the most significant factors. Both were associated with an increased probability to reach reproduction, potentially linked to poaching. Degree of inbreeding was negatively associated with the probability to reach reproduction, while gravel road density and being born in Sweden were positively associated with it. Our findings suggest an influence of legal and illegal human activities on the juvenile stage of wolves for the probability to reach reproduction. Our study enhances the understanding of how early-life conditions and intrinsic traits shape reproduction and underscores the challenges of wolf conservation in anthropized landscapes.

## Introduction

Life-history theory seeks to explain the general features of an organism’s life cycle, including both intra- and interspecific variation, by exploring how organisms allocate resources to growth, survival, and reproduction throughout their lives [1,2]. These resource allocation strategies shape their overall evolutionary patterns. A key component of an organism’s life-history strategy is its ability to reach reproductive maturity. To do this, most individuals must survive through distinct life stages such as the juvenile stage followed by the adult stage, where they can find a mate and reproduce. Success through these stages depends on an individual’s fitness, which is influenced by intrinsic traits and environmental factors.

The juvenile stage is particularly crucial as it is often characterized by low and variable survival rate stages [2,3], with early-life environmental conditions having lasting effects on an organism’s biology. Indeed, favourable early-life conditions can enhance an individual’s chances of survival and increase the probability of reaching reproductive maturity, whereas adverse early life conditions can lead to reduced body growth, affect behaviour or physiological processes, such as delaying reproduction, and reduce overall fitness [4–7]. Anthropogenic habitats affect fitness and life history traits of several species in various ways. For instance, [8] show that modified habitats negatively affect the fitness by modifying the life history traits, favouring a faster pace-of-life with earlier dispersal and dominance acquisition in a cooperatively breeding species. [9] highlight how individuals behaviourally respond to various types of human disturbances. These responses can be direct or indirect, often resulting from changes in factors that affect fitness, such as resource availability, opportunities and success in dispersal, access to free space, and the presence of other interacting species.

Due to their large home ranges and long generation times [10] resulting in slow population recovery, large carnivores might be especially vulnerable to human-induced changes such as the fragmentation and alteration of their habitat [11]. Over the past few decades, large carnivores have done a notable recovery, establishing in anthropized landscapes [12]. This can lead to conservation conflicts, particularly in rural areas where livestock depredation becomes a pressing concern for farmers and herders [12–15]. In addition, as many large carnivore populations remain small and isolated, they are more exposed to threats affecting the long-term viability of populations such as loss of genetic variation, inbreeding depression and reduced adaptive potential [16–18].

One of the most remarkable large carnivore recoveries in Europe is that of the wolves (*Canis lupus*). As highly adaptable species, wolves are able to settle along the entire gradient from low to high human impact [12]. Cohabitation with humans and environmental factors impact both wolf behaviour and population dynamics [19,20]. For instance, roads pose a trade-off for wolves, with negative effects linked to humans [21] including disturbance [22], increased traffic mortality [23] and increased access for poachers [24]. Positive effects may result from increased ease of travel [25–27], efficient scent-marking [28] and access to prey [29,30]. Similarly, human density can also negatively impact the fitness of wolves as they tend to avoid human infrastructure [22,31], in particular for den and rendezvous sites [32]. The quality and quantity of resources available during early life is one of the well-known factors affecting the fitness and performance later in life [33]. Besides prey density, the quantity of food available for consumption is also contingent upon the wolves’ hunting success. In turn, hunting success [34–36] and kill rate [37,38] of wolves may be affected by climatic conditions such as snow depth. Resource availability also depends on intra-specific competition as high wolf densities lead to reduced territory size [20,39]. On the other hand, higher wolf densities might also have a positive impact on the probability to reach reproduction by increasing the chances to find a mate, which can be challenging at low population densities [14,40–42].

Intrinsic traits can also affect the fate of juveniles. Inbreeding, known for its detrimental effects across many species [43–46] including wolves [47–50], has been shown to have a significant impact on juveniles [43,51]. Inbreeding depression can lead to malformations [48,52,53], increase the age of first reproduction [42], as well as decrease the pairing and breeding success [50]. Furthermore, the probability to reach reproduction can differ among sexes, with juvenile male mammals commonly exhibiting higher mortality rates than their female counterparts [54–56]. While radio tagging animals can give insight into intrinsic characteristics, the tags or collars themselves can affect the behaviour, survival and reproduction of the studied individual. Although the negative impact of GPS-collars on bird survival and reproduction is well-documented [57], their effect on mammals remains less well studied. Studies on collared large carnivores have presented different conclusions where some report a higher survival of collared wolverines (*Gulo gulo*) [58] and wolves [59,60] whereas others link collars to a higher risk of mortality for wolves [61,62].

The recovery of the Scandinavian wolf population serves as a well-documented example of the re-establishment of wolves in Europe. After the first reproduction as early as 1983, only five immigrating individuals contributed with unique haplotypes during the 1983-2008 period [50], and with wolf numbers estimated to 375 (CI: 352 to 402) wolves in Scandinavia in 2019 [63]. Compared to other regions worldwide, the population was still at a relatively low density with a high ratio of moose (*Alces alces*) to wolf [30,64], with moose being the primary prey species in this population [65–67]. Due to extensive monitoring efforts of the population, which has allowed for the identification of 97% of all reproductive events, the natal territory and the inbreeding level is known for almost all individuals from the re-establishment of the population in 1983 to date [68]. The population also serves as a good example of the conservation challenges linked to wolf recovery as the historical bottleneck during recolonization has led to severe inbreeding depression [49]. [50] showed that offspring from immigrants have higher pairing and breeding success compared to offspring from native and more inbred pairs. Similar to other regions worldwide, the recovery of wolves in Scandinavia has resulted in conflicts within local communities [13,14] that pose challenges for the conservation of the wolf population. Poaching has been estimated to account for half of the total wolf mortality [69]. Beyond poaching, wolves have been legally culled in Scandinavia for damage control and through quota systems [70]. However, management goals and wolf policy differ between Sweden and Norway, e.g. in population size and distribution [71,72]. In Sweden, wolves are allowed to settle outside the reindeer husbandry area (approximately 55% of the total country area of 447 425 km²), and in Norway, wolves are allowed to settle in a ‘wolf zone’ (approximately 5% of the total country area of 324 220 km^2^). Norway is also characterised by a generally lower social acceptance of large carnivores as compared to Sweden [13].

After the juvenile stage, wolves can disperse or stay philopatric before settling in a territory with a mate and reproduce. In this study, we address specifically the juvenile life history stage of wolves and its impact on later fitness until first reproduction in the wolf population of Scandinavia. We defined the probability of reaching reproduction as the probability of reproducing and having pups surviving at least five months of age. To better understand how the conditions in the juvenile stage can affect the probability to reach reproduction, we examined the influence of extrinsic factors related to the environment of the natal territory and intrinsic factors such as inbreeding, sex, and collaring in the Scandinavian wolf population. Our hypotheses are summarized in Table 1.

**Table 1.**
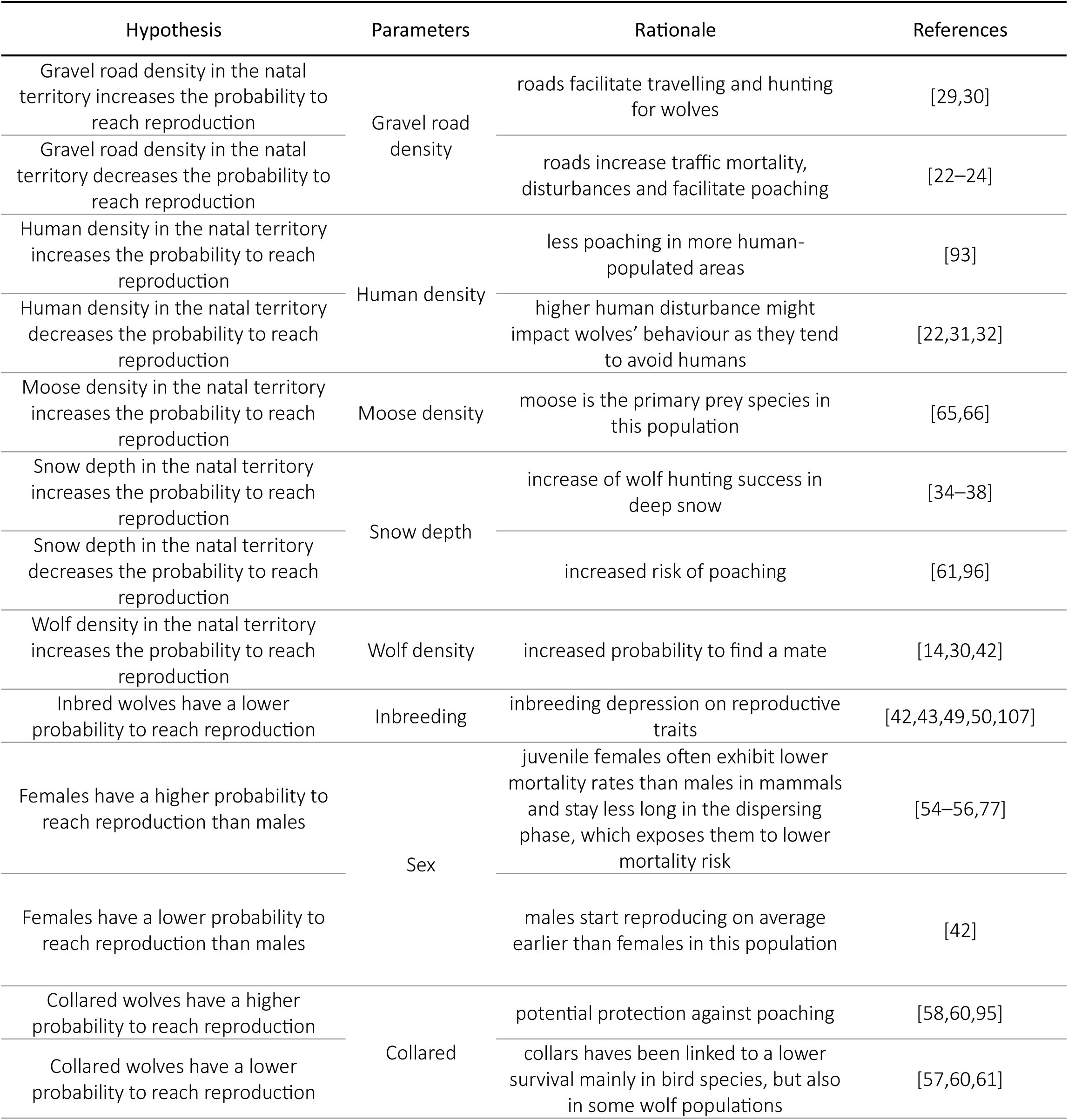
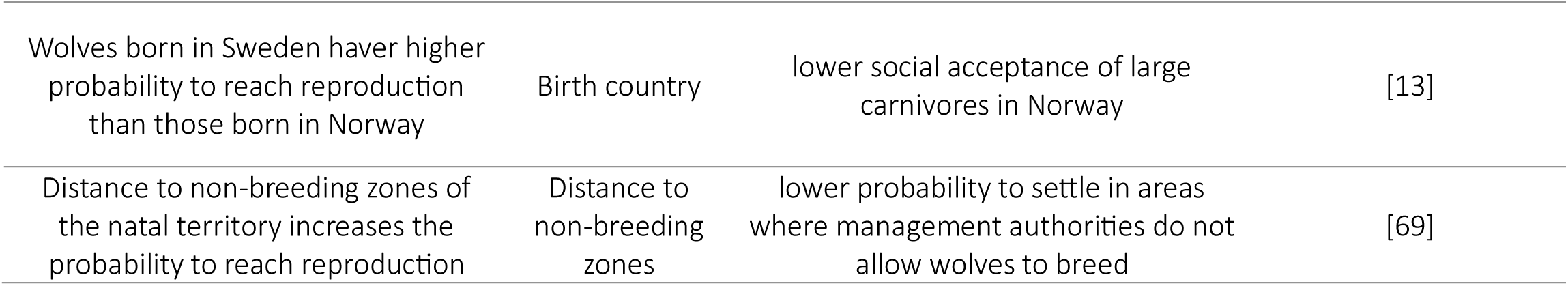
Hypotheses on the effects of extrinsic factors (gravel road density, human density, moose density, snow depth, wolf density, distance to non-breeding zones and birth country) and intrinsic factors (inbreeding, sex, being collared) on the probability for wolves to reach reproduction in Scandinavia.

| Hypothesis | Parameters | Rationale | References |
| --- | --- | --- | --- |
| Gravel road density in the natal territory increases the probability to reach reproduction | Gravel road density | roads facilitate travelling and hunting for wolves | [29,30] |
| Gravel road density in the natal territory decreases the probability to reach reproduction |  | roads increase traffic mortality, disturbances and facilitate poaching | [22–24] |
| Human density in the natal territory increases the probability to reach reproduction | Human density | less poaching in more human-populated areas | [93] |
| Human density in the natal territory decreases the probability to reach reproduction |  | higher human disturbance might impact wolves' behaviour as they tend to avoid humans | [22,31,32] |
| Moose density in the natal territory increases the probability to reach reproduction | Moose density | moose is the primary prey species in this population | [65,66] |
| Snow depth in the natal territory increases the probability to reach reproduction | Snow depth | increase of wolf hunting success in deep snow | [34–38] |
| Snow depth in the natal territory decreases the probability to reach reproduction |  | increased risk of poaching | [61,96] |
| Wolf density in the natal territory increases the probability to reach reproduction | Wolf density | increased probability to find a mate | [14,30,42] |
| Inbred wolves have a lower probability to reach reproduction | Inbreeding | inbreeding depression on reproductive traits | [42,43,49,50,107] |
| Females have a higher probability to reach reproduction than males | Sex | juvenile females often exhibit lower mortality rates than males in mammals and stay less long in the dispersing phase, which exposes them to lower mortality risk | [54–56,77] |
| Females have a lower probability to reach reproduction than males |  | males start reproducing on average earlier than females in this population | [42] |
| Collared wolves have a higher probability to reach reproduction | Collared | potential protection against poaching | [58,60,95] |
| Collared wolves have a lower probability to reach reproduction |  | collars have been linked to a lower survival mainly in bird species, but also in some wolf populations | [57,60,61] |
| Wolves born in Sweden have higher probability to reach reproduction than those born in Norway | Birth country | lower social acceptance of large carnivores in Norway | [13] |
| Distance to non-breeding zones of the natal territory increases the probability to reach reproduction | Distance to non-breeding zones | lower probability to settle in areas where management authorities do not allow wolves to breed | [69] |

## Materials & Methods

### I. The wolf population on the Scandinavian Peninsula

The studied wolf population is distributed in south-central Sweden and in the adjacent areas in south-east Norway. Wolves are prevented from settling as residents in areas where reindeer husbandry occurs [73]. In Norway, the wolf zone represents the only area where wolves are allowed to reproduce (Figure 1).

**Figure 1.**
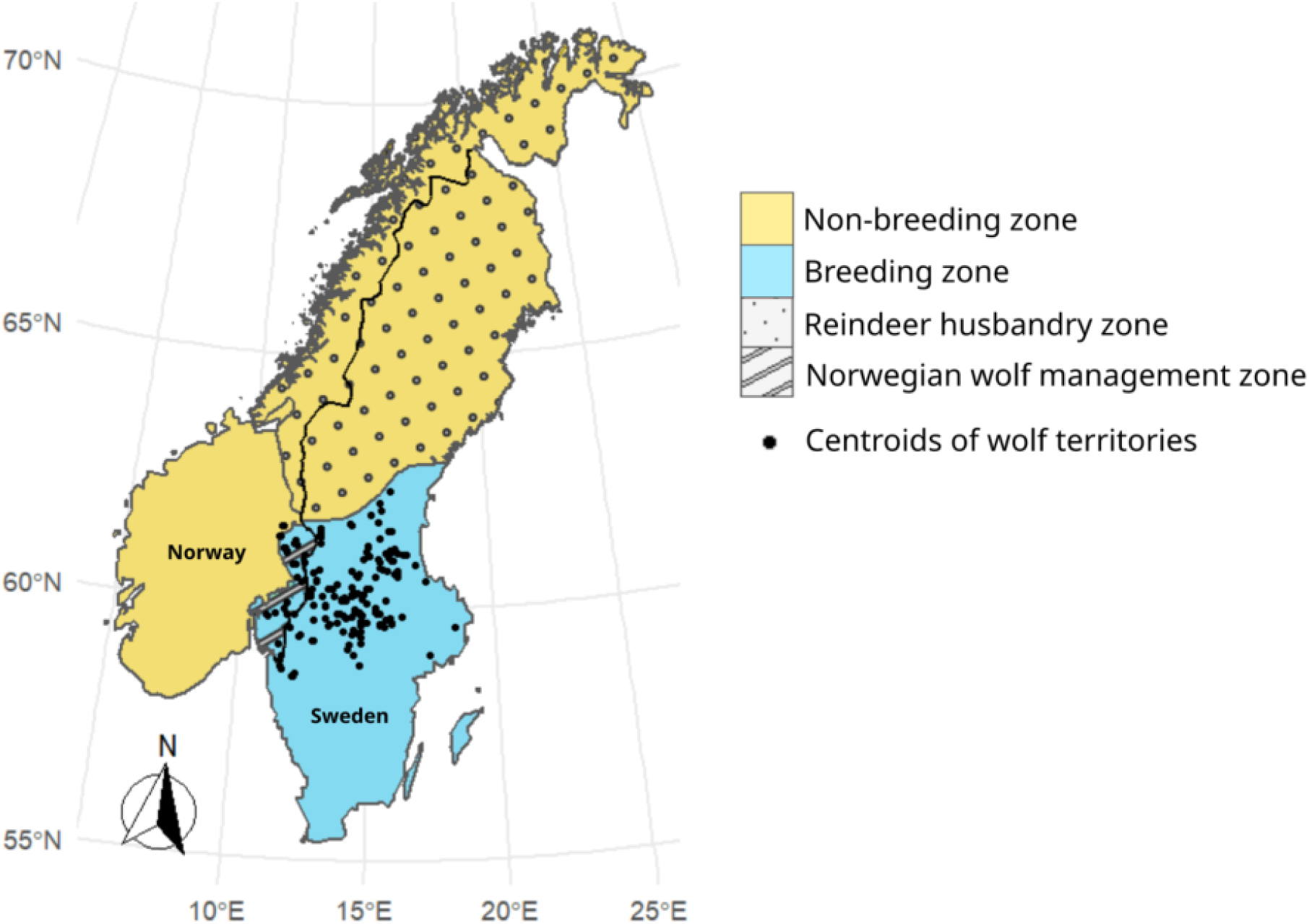
Map of the study area with the Norwegian wolf management zone and the Scandinavian reindeer husbandry zone. The blue part represents the area where wolves are legally accepted to settle and reproduce (the breeding zone) and the yellow one is the non-breeding zone. The black dots represent the centroids of the wolves’ natal territories used in this study from 2003 to 2016.

After being declared functionally extinct in 1966, the wolf population has re-established on the Scandinavian peninsula in the late 1970s with the immigration of few wolves coming from the Finnish– Russian population [14,49,74]. The population has been increasing, reaching approximately 460 (CI: 439 - 483) wolves in Scandinavia in the beginning of the monitoring season 2020-2021 with less than 20% of the wolves ranging in Norway and the rest in Sweden [75].

Every year since 1998, a monitoring programme has been conducted during the winter period (October 1^st^ -March 31^st^) for individual identification, sex, and parentage analysis. It was originally based only on snow tracking, but from the early 2000’s it also included DNA analyses of non-invasive samples (scat, urine, hair). Based on these data, territorial pairs and packs can be identified in order to determine the annual number of reproduction events, confirmed as described by [68]. The DNA analyses for individual identification and relatedness enable the reconstruction and annual update of the pedigree of the population, which provides annual estimates of inbreeding of almost all individuals in the Scandinavian population [49,68].

### II. Selection of target wolf individuals

The identification of wolf individuals in this study was based on DNA sampling from i) scats, urine, or oestrus blood collected during the monitoring season, ii) saliva from depredation events, or iii) blood sampled from alive captured wolves. This study utilized data from 2003 to 2021, including individuals born in Scandinavia from 2003 to 2016. A previous study has shown that 95% of the surviving wolves had reproduced by the age of five years and that the median age at first reproduction in this part of the population was three years for females and two years for males [42]. Thus, to minimize the risk of inaccurately classifying wolves as non-breeding when they have the potential for reproduction, our study included only wolves born up to 2016.

To minimize bias towards individuals that successfully bred and to maintain consistency in our data, our study included only those individuals that were identified as being alive during minimum their first monitoring season [68], which spans from 5 to 11 months of age (1^st^ October – 30^th^ March). This approach was taken because older individuals exhibit a higher probability of reaching reproduction. Our sample included 340 individuals for which the birth year was known, either because they were sampled and individually identified during the first year after their parents’ first reproduction event (n = 310) or they were captured and identified as pups (<1 year old, n = 30). Additionally, 242 individuals with an unknown age were included in the study that resulted in a total dataset of 582 (Table 2), provided their first identification was within their natal territory at during the monitoring season. It is likely that the majority of these individuals were less than one year old, i.e. identified during the first monitoring season after birth, as 76% of the pups permanently leave their natal territory before their second monitoring season, i.e. before the age of 1 year and 5 months [76,77].

**Table 2.** Number of individuals in the Scandinavian wolf population that reached reproduction in the different datasets (2003-2021).

|  | Total number of individuals | Number of individuals<br>that reached<br>reproduction |
| --- | --- | --- |
| Excluding legally killed<br>(main analysis) | 456 | 108 (23.6%) |
| Including legally killed<br>(Appendix 1) | 582 | 122 (20.9%) |
| Excluding collared<br>individuals<br>(Appendix 2) | 437 | 99 (22.6%) |

Across the whole dataset of 582 wolves, 126 had been legally killed (Table 2). Because the decision to kill a wolf can depend on factors beyond our research scope, we excluded those wolves that were legally killed from the main analysis and we conducted a separate analysis including the legally killed individuals (Appendix 1). To ensure consistency, we applied the same threshold of five years of age for legally killed wolves, regardless of the fact that they breed or not, i.e. if a wolf was killed before reaching five years of age, we assumed it may not have had the time to breed yet and therefore was removed from our analysis.

### III. Intrinsic factors

The intrinsic factors related to individual wolves were inbreeding coefficient, sex, and being fitted with a collar. The inbreeding coefficient of each individual is based on the pedigree of the Scandinavian wolf population [49,78]. As the objective was to study the effect of collaring during the early life stage of wolves, we considered as collared individuals only those wolves collared before 1 year of age (n=19 excluding legally killed individuals and n=27 including them). We also conducted a separate analysis excluding these collared wolves (Appendix 2, Table 2).

### IV. Extrinsic factors

The extrinsic factors consisted of environmental and anthropogenic variables linked to the natal territory. The measure of natal territory used to extract data was a circular buffer around the centroid of the natal territory polygon as registered during the annual monitoring. The size of the natal territory created was 1000 km^2^ (18 km radius buffer), as represented by the average size of a Scandinavian wolf territory [79].

Human density was calculated as the yearly number of inhabitants per km^2^ for each municipality for Sweden (https://www.scb.se/) and Norway (https://www.ssb.no/). For each estimated natal territory, human density was measured as the average human density of the municipalities overlapping with the natal territory. Gravel road density was calculated as the average length of gravel roads (km per km^2^). Wolf density was estimated as the number of bordering neighbouring territories, i.e. the number of territories overlapping with the natal territory. As moose harvest size has shown to be correlated with the population density of moose in Scandinavia [80], we used the yearly hunting bag records as an index of moose density (number of moose killed/10 km^2^ for counties in Sweden www.algdata.se and Norway https://www.ssb.no/). Data on hunting bag records was generated as a weighted average of the moose density of the counties overlapping with the territory. Yearly average of snow depth was estimated from daily snow depth data extracted from the database SMHI for the weather stations in Sweden (https://www.smhi.se) and from website seklima (https://seklima.met.no/.) using the data from met.no (https://www.met.no) for Norway. Corrections for missing values (3.3 % in Sweden and 11.7% in Norway) were implemented according to the SMHI recommendations (Appendix 3). As we were mainly interested in the effect of snow depth during the first year of life of wolves, we used the yearly average of snow depth data from 1^st^ of May (average birth of pups) to 30^th^ of April next year, excluding the summer months (July, August, September), for each weather station. Consequently, for each individual, the average snow depth during its first year of life in its natal territory was estimated using a kriging interpolation model including the effect of altitude as this factor showed to improve the model accuracy [82]. We further included two geographical factors of the natal territory, i.e. the country of birth (Norway or Sweden according to the location of the territory centroid), as well as the distance between the natal territory centroid and the closest area where the wolves were not allowed to establish (non-breeding zone), i.e. outside of the Norwegian wolf zone and inside the reindeer husbandry area (Figure 1).

### V. Statistical analysis

We defined whether an individual has reached reproduction or not as a binary response variable in our model, assigning a value of 1 for individuals which had pups confirmed alive during the following monitoring season and 0 for those without. We employed a generalized linear mixed-effects model (GLMM) using the glmmTMB R package version 1.9.14 [83], with a binomial distribution (logit link) that included ten fixed effects (sex, country, and collared as binary variables, while inbreeding, human density, gravel road density, snow depth, moose density, distance to non-breeding zones for wolves, and wolf density as scaled continuous variables) and the ID of the parental pair included as a random effect to account for potential correlations within the data, as offspring from the same parental pair may share genetic or environmental characteristics that could introduce non-independence among observations.

The correlation between the explanatory variables was assessed by a correlation test (cor.test function in R) and by calculating the variance inflation factor (check_collinearity function from the R package performance version 0.11.0 [84]). None of the explanatory variables had to be excluded due to collinearity (all pairwise Pearson r < 0.39) and the highest variance inflation factor was 2.77 for country of birth. No deviation from the model assumptions were detected in the full model using the simulateResiduals function from the DHARMa package in R version 0.4.6 [85]. As we included a large number of variables, we used a model selection procedure to determine the combination of variables that best fit the data. Model selection was performed using the dredge function from the MuMIn package version 1.47.5 [86], based on the Akaike Information Criterion corrected for small sample sizes (AICc). To ensure robustness, only models with a ΔAICc value of 2 or less were retained for model averaging with the function model.avg of the MuMIn package.

## Results

Among the 456 individuals identified alive as pups throughout their first monitoring winter, 108 (24%) reproduced with at least one pup confirmed alive during the following monitoring season. Including also wolves that were legally killed, the probability to reach reproduction was reduced to 21% (Table 2).

In our analysis, 15 models had ΔAICc ≤ 2 and were included in the model averaging. Two factors, human density, and being collared, were present in all of the best models after model selection (Table 3). The confidence intervals of these factors in the averaged model did not overlap with zero (Figure 2).

**Figure 2.**
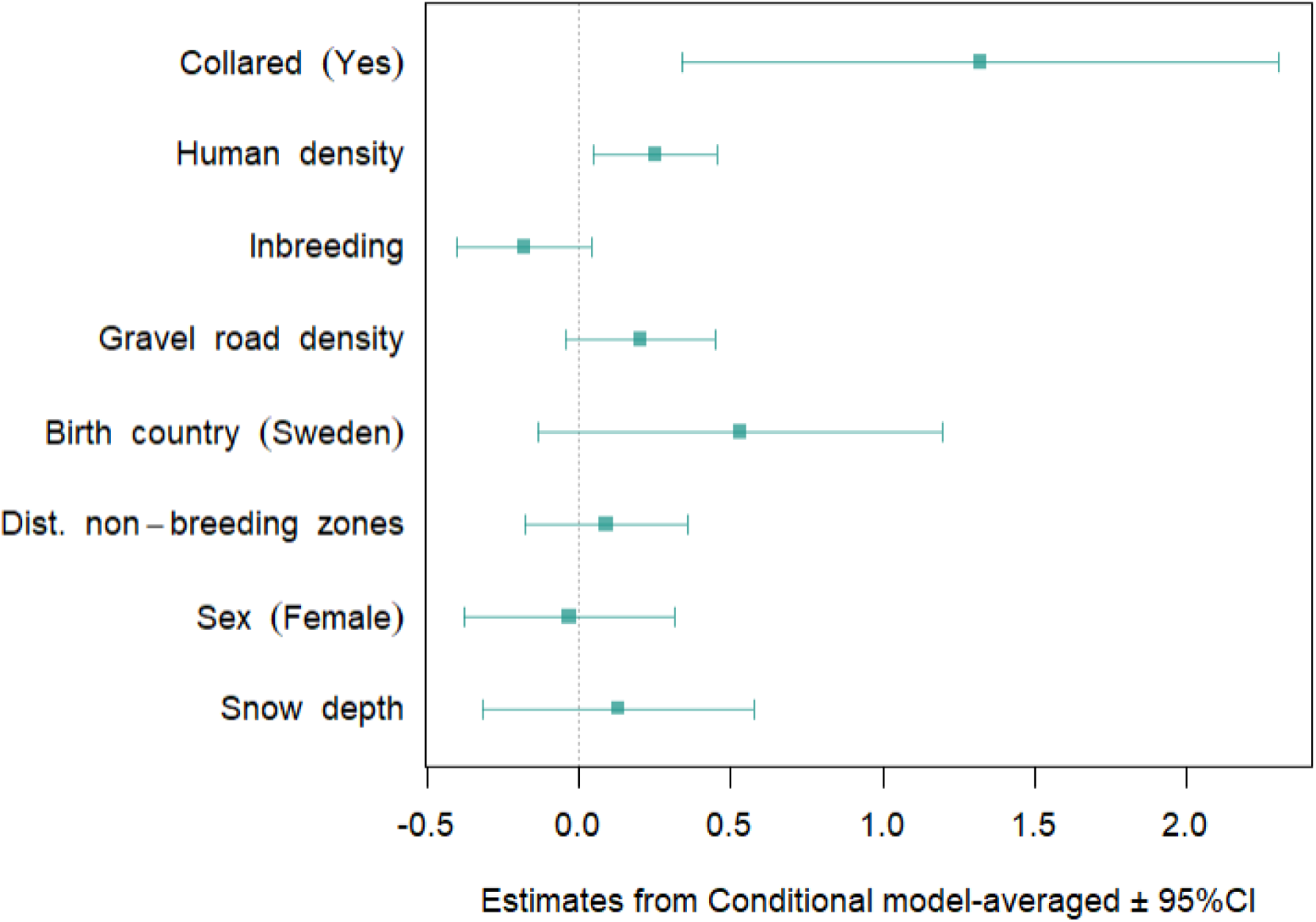
Coefficient plot of the averaged model of factors impacting the probability of reaching reproduction for wolves in Scandinavia. The bars represent the 95% confidence interval of the coefficients estimated by the averaged model (Table 3).

**Table 3.** Model selection and model-averaged parameter estimates (β), standard error (SE) for conditional and full averaged model on the probability to reach reproduction for wolves in Scandinavia during 2003-2021. The models were ranked based on AICc and only the top models (ΔAICc ≤ 2) were kept and used for model averaging. The best model has an AICc of 495.05. The reference in the analyses is “No” for the predictor Collared, “Norway” for Country and “Male” for Sex. *** < 0.001 < ** < 0.01 < * < 0.05

|  |  | Intercept | Collared | Human density | Inbreeding | Gravel road density | Birth country | Dist. non-breeding zones | Sex | Snow depth | df | ΔAICc | weight |
| --- | --- | --- | --- | --- | --- | --- | --- | --- | --- | --- | --- | --- | --- |
|  |  | X | X | X | X | X | X |  |  |  | 7 | 0 | 0.12 |
|  |  | X | X | X |  | X | X |  |  |  | 6 | 0.23 | 0.11 |
|  |  | X | X | X | X |  | X |  |  |  | 6 | 0.55 | 0.09 |
|  |  | X | X | X | X |  |  |  |  |  | 5 | 0.75 | 0.08 |
|  |  | X | X | X | X | X |  |  |  |  | 6 | 0.88 | 0.08 |
|  |  | X | X | X |  | X |  |  |  |  | 5 | 1.30 | 0.06 |
|  |  | X | X | X |  |  | X |  |  |  | 5 | 1.38 | 0.06 |
|  |  | X | X | X | X | X | X |  |  | X | 8 | 1.54 | 0.06 |
|  |  | X | X | X | X | X | X | X |  |  | 8 | 1.61 | 0.05 |
|  |  | X | X | X |  |  |  |  |  |  | 4 | 1.65 | 0.05 |
|  |  | X | X | X | X | X | X |  | X |  | 8 | 1.73 | 0.05 |
|  |  | X | X | X | X |  |  | X |  |  | 6 | 1.77 | 0.05 |
|  |  | X | X | X |  | X | X |  |  | X | 7 | 1.96 | 0.05 |
|  |  | X | X | X |  | X | X | X |  |  | 7 | 1.96 | 0.05 |
|  |  | X | X | X |  | X | X |  | X |  | 7 | 1.99 | 0.04 |
| Full model | β | -1.55*** | 1.32** | 0.25* | -0.10 | 0.13 | 0.36 | -5x10<sup>-3</sup> | 0.01 | 9x10<sup>-3</sup> |  |  |  |
|  | SE | 0.33 | 0.50 | 0.10 | 0.12 | 0.14 | 0.37 | 0.07 | 0.08 | 0.05 |  |  |  |
| Conditional averaged | β | -1.55*** | 1.32** | 0.25* | -0.18 | 0.20 | 0.53 | -0.03 | 0.13 | 0.09 |  |  |  |
|  | SE | 0.33 | 0.50 | 0.10 | 0.11 | 0.12 | 0.34 | 0.18 | 0.23 | 0.14 |  |  |  |

The probability to reach reproduction was positively associated with human density in the natal territory, with an increase of one standard deviation in the number of inhabitants/km^2^ resulting in a 1.28 times higher odds (exp(0.25), Table 3) of reaching reproduction. To illustrate this point, the average predicted probability of reproduction was 16% [CI = 7%-25%] in natal wolf territories with a human density of 1 inhabitant/km^2^, and 27% [CI = 13%-41%] in territories with 100 inhabitants/km^2^ (Figure 3).

**Figure 3.**
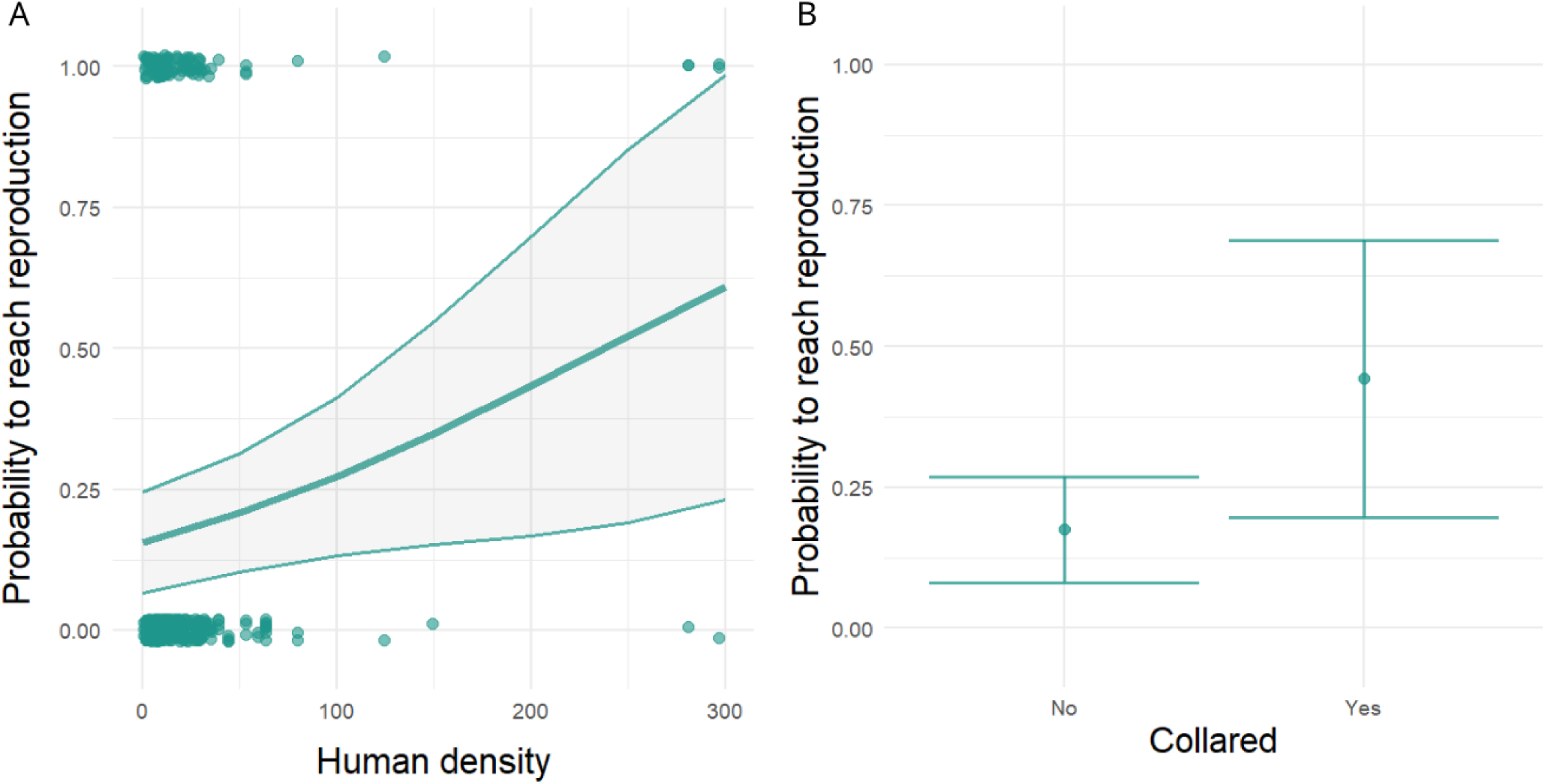
Probability to reach reproduction in relation to A) the human density in the natal territory (inhabitant/km^2^) and B) being collared. The lines indicate the fitted values, with associated 95% confidence interval from the model-averaged estimates (Table 3). For panel A, sex was held constant at “Male”, birth country at “Norway”, collared at “No”, and the other continuous variables coefficients (inbreeding, human density, gravel road density, snow depth, moose density, distance to non-breeding zones for wolves, and wolf density) were kept at their mean. The dots in the panel A correspond to the observed data points, where their position on the y-axis represents the actual binary outcome of reaching reproduction (0 or 1).

Among the 19 wolves collared during their first year of life, reproduction was recorded for 9 individuals, resulting in a probability to reach reproduction of 44% [CI = 20%-69%] as compared to 17% [CI = 8%-27%] (Figure 3) for non-collared individuals. Collared wolves had more than three times higher odds (odds ratio = 3.74) of reaching reproduction than non-collared wolves (Table 3).

The density of gravel roads in the natal territory and the country of birth were both retained in ten models, among the 15 best models. Both wolves born in Sweden or in a natal territory with higher road density were associated with a higher probability to reach reproduction (Table 3, Appendix 4). Inbreeding was retained in eight of the 15 best models and showed a negative but non-significant trend on the probability to reach reproduction (Table 3, Appendix 4). The other explanatory factors (distance to non-breeding zones, sex and snow depth) were retained in only a few of the top-rated models except moose and wolf density that were not included in any of the top-rated models (Table 3).

The inclusion of legally killed individuals (Appendix 1) or the removal of collared wolves (Appendix 2) gave similar results and did not change the main outcomes. In our data, 55% of the individuals were killed legally within the area where wolves are allowed to reproduce, and 64% of these events took place in Sweden, which also accounts for 78% of the individuals in our study.

## Discussion

We identified two relevant indirect drivers related to the probability of reaching reproduction for wolves, including human density within the natal territory and whether the wolf was fitted with a collar during its first year of life. Moreover, we observed weak evidence for gravel road densities within the natal territory to be positively related to the probability of reaching first reproduction. There was also weak evidence for higher probability to reach first reproduction if wolves were born in Sweden rather than Norway. Finally, in line with our hypothesis, we observed a tendency for a higher probability of reaching first reproduction for less inbred wolves. Contrary to our hypotheses, we did not find support for an effect of the other predictors, i.e. moose density, wolf density, sex, snow depth, and the distance to non-breeding wolf zones.

The strong relation of reaching reproduction with human density and collaring may reflect complex and indirect interactions between wolves and human activities, including poaching of wolves. Even though the major consensus existing in many studies is that large social carnivores like wolves are conflict prone in areas with high human densities [22,32,87,88], several studies have also suggested that wolves are highly capable of persisting in human-altered landscapes [89–91]. A higher abundance of ungulate prey in agricultural areas [92] has been suggested as a potential reason for the selection of human-altered habitats over natural ones by red wolves (*Canis rufus*). In our study, the observed positive relation between probability of reaching reproduction and human density in the natal territory could be functionally linked to easier access to prey in more inhabited and agricultural areas. Another potential explanation is that higher human densities may discourage poaching due to an increased risk of being discovered and caught by legal enforcement [93]. Relatedly, the acceptance of large carnivores, including wolves, tends to be higher in more urbanized areas, whereas inhabitants in rural areas generally express a more negative attitude [13,94].

The fact that wolves collared within their first year exhibited a substantial advantage in the probability of reaching reproduction supports the finding presented by [58]. That study showed indeed a positive effect of GPS-collars on the survival of wolverines in Scandinavia and the authors attributed these results to the collars acting as a deterrent against illegal killing [58]. An alternative explanation for our finding could be that wolves, learning from past capture experiences, develop an avoidance behaviour towards humans, by perceiving these events as traumatic. If this avoidance behaviour is realized, it could make wolves less exposed and vulnerable to human presence and activities, including poaching. If true, this finding also gives rise to several inquiries regarding the validity of utilizing collared individuals to estimate poaching rates, a common approach used for many studies [61,69,70,93,95,96]. Therefore, the estimation of mortality rate in general and poaching rate in particular and its consequences in the Scandinavian population based on collared individuals [69,70] might have been underestimated. A third explanation could be a bias in our sample of collared wolves as these generally are captured in the latter half of the monitoring season whereas DNA-sampling of non-collared wolves may be done during the first half of the monitoring season. In this scenario, collared wolves may be those that had a higher survival as compared to those that survived only to their first sampling event. Nonetheless, considering that our results have been obtained from a limited number of collared individuals (5% of the total sample), prudence is advised when interpreting the implications. Future research should indeed examine more in depth the fitness consequences of fitting wolves with collars by comparing the proportion of collared individuals with the uncollared segment of the population.

The observed tendency for a negative impact of inbreeding on the probability of reaching reproduction aligns with the important role of genetic diversity for population growth and persistence [17]. Although with lower effect size, our study in line with the previous findings on negative impact of inbreeding on fitness of the Scandinavian wolf population [42,49,50,52,53]. The low effect size may have several underlying causes. For one reproducing immigrant, the pack dissolved for unknown reasons and none of the offspring reproduced. In another case, the human translocation of a pair, where both parents were immigrants, may have affected the reproductive probability of their offspring and none were successfully reproducing. Another potential reason of the weak relation with inbreeding may be related to the interplay of inbreeding with environmental and anthropogenic stressors characterizing the Scandinavian wolf population. Inbreeding often interacts with the environment leading to a stronger disadvantage of inbred individuals in stressful environments [97]. Studies conducted in less stressful environments, characterized by for example more food sources and less competition, may provide weaker results and therefore lack statistical power to detect the true effects of inbreeding depression [97]. In the Scandinavian wolf population, where poaching has posed a greater threat than natural causes [69,70], inbred individuals may experience less pronounced effects, as the impact of poaching potentially could overshadow the consequences of inbreeding at the population level.

The higher tendency of reaching reproduction for wolves born in Sweden as compared to Norway aligns with our hypothesis, which is grounded in the presumed lower social and legal acceptance of wolves in Norway [13]. Although there is only one wolf population on the Scandinavian peninsula, the conflicts surrounding wolves and their management are different for Sweden and Norway, even though they share many social, cultural, and geographical characteristics [98]. Norway has a smaller geographical area where wolves are allowed to breed (5% compared to 55% for Sweden) as well as a lower maximum population size threshold before the population is limited by human control [99]. However, related to population size, our data did not indicate a difference in the distribution of culling rates between the two countries. Besides these differences in management strategy, the poaching of large carnivores including wolves was more accepted in Norway than in Sweden in a study conducted before 2013 [13]. In fact, Sweden can be described as a source for the Norwegian part of the population and Norway is reliant on Sweden to meet its management objectives for wolf conservation [71,100]. Contrary to previous results [72] and to our hypothesis, the distance between the natal territory and non-breeding zones showed no discernible impact on the probability to reach reproduction.

The tendency for a positive relation of the probability of reaching reproduction with gravel road density could fit our hypothesis of gravel roads facilitating wolf movement and predation opportunities. Higher densities of gravel roads may lead to lower energy requirement for movement, which could positively affect wolf body condition before dispersal, which may in turn increase the probability of finding a territory as well as a mate and reproduce. This potential explanation is in contrast with the alternative hypothesis that higher densities of gravel road may facilitate poaching and consequently reduce the probability of reaching reproduction for wolves. In addition, the lack of correlation between the probability of reaching reproduction and moose density may be explained by the high density of moose or alternative ungulate prey across the distribution range of wolves in Scandinavia [64,79,101], which may therefore not represent a limiting factor for wolf fitness. Similarly, the lack of support for an effect of sex on the probability of reaching reproduction may be related to the monogamous nature of wolves with virtually no divorces [102].

Human activity is recognized to exert various influences on wolf populations in Europe, by impacting their behaviour [25,31], their population dynamics [70], genetics [103] and distribution [104,105]. In Scandinavia, the conflict with humans poses significant challenges for the conservation of the wolf population as poaching has been estimated to account for up to half of the total wolf mortality and severely limits population growth [69]. While our results suggest that human density in the natal territory and collaring within the first year of life have a positive relationship with the probability of reaching reproduction for wolves, further research is warranted to disentangle the mechanisms driving such associations. This should possibly include the environment experienced in later life-stages but preceding first reproduction, i.e. dispersal and territory establishment. Indeed, wolf individuals can either be philopatric and remain in the natal habitat or disperse to new environments, a decision that can have an impact on their chance to survive and find a mate, finally affecting their probability to reach reproduction. Overall, with a focus on the juvenile early-life stage, our findings contribute to our understanding of how early-life conditions experienced in the natal territory and intrinsic traits affect wolves’ probability of reaching reproduction but also highlight the conservation challenges of wolves coexisting with humans in increasingly anthropized landscapes.

## Ethics

All procedures including capture, handling and GPS collaring of wolves fulfilled ethical requirements and have been approved by the Swedish Animal Experiment Ethics Board (permit no. C 281/6) and the Norwegian Experimental Animal Ethics Committee (permit no. 2014/284738-1). The GPS data were collected into the Wireless Remote Animal Monitoring database system for data validation and management [106].

## Data accessibility

Data will be available from the Dryad Digital Repository upon acceptance.

## Author’s contributions

Funding acquisition: C.W., M.Å., H.S.; Concept: L.A., C.D.B., C.W., M.Å., H.S.; Data compilation: M.Å., Ø.F., H.S., C.W., L.A., C.D.B.; Statistical analyses and figures: L.A.; Manuscript writing: L.A., C.D.B.; Manuscript commenting: All authors.

## Acknowledgements

We thank all the people who conducted fieldwork, laboratory work, and/or administered the monitoring of wolves in Sweden and Norway during the last decades. We also thank the capture crew, especially J.M. Arnemo, P. Ahlqvist, U. Grinde, T.H. Strømseth, A.L. Evans, D. Ahlqvist, and B. Fuchs, for capturing and handling the wolves.

## Supporting Information

**Appendix 1.** Model selection and model-averaged parameter estimates (β), standard error (SE) and for conditional and full averaged model including legally killed individuals. The models were ranked based on AICc and only the top models (ΔAICc ≤ 2) were kept and used for model averaging. The best model has an AICc of 593.86. The reference in the analyses is “No” for the predictor Collared, “Norway” for Country and “Male” for Sex. *** < 0.001 < ** < 0.01 < * < 0.05

| | | Int. | Collared | Human density | Inbreeding | Gravel road density | Country | Wolf density | Dist non-breeding zones | Sex | Snow depth | Moose density | df | $\Delta AICc$ | weight |
| --- | --- | --- | --- | --- | --- | --- | --- | --- | --- | --- | --- | --- | --- | --- | --- |
|  |  | X | X | X | X |  |  |  |  |  |  |  | 5 | 0 | 0.13 |
|  |  | X | X | X | X | X |  |  |  |  |  |  | 6 | 0.24 | 0.12 |
|  |  | X | X | X | X | X | X |  |  |  |  |  | 4 | 1.14 | 0.08 |
|  |  | X | X | X |  | X |  |  |  |  |  |  | 5 | 1.15 | 0.07 |
|  |  | X | X | X | X |  | X |  |  |  |  |  | 6 | 1.16 | 0.07 |
|  |  | X | X | X | X |  |  |  |  | X |  |  | 6 | 1.37 | 0.07 |
|  |  | X | X | X |  |  |  |  |  |  |  |  | 7 | 1.41 | 0.07 |
|  |  | X | X | X | X | X |  |  |  | X |  |  | 7 | 1.47 | 0.06 |
|  |  | X | X | X | X |  |  | X |  |  |  |  | 6 | 1.64 | 0.06 |
|  |  | X | X | X | X |  |  |  |  |  |  | X | 6 | 1.71 | 0.06 |
|  |  | X | X | X | X |  |  |  | X |  |  |  | 6 | 1.77 | 0.06 |
|  |  | X | X | X | X |  |  |  |  |  | X |  | 6 | 1.79 | 0.05 |
|  |  | X | X | X | X | X |  |  |  |  |  | X | 6 | 1.85 | 0.05 |
|  |  | X | X | X |  | X | X |  |  |  |  |  | 5 | 1.96 | 0.05 |
| Full model averaged | $\beta$ | -1.46*** | 0.95 * | 0.24* | -0.15 | 0.07 | 0.05 | 4x10<sup>-3</sup> | 3x10<sup>-3</sup> | 0.02 | -3x10<sup>-3</sup> | 7x10<sup>-3</sup> | | | |
|  | SE | 0.17 | 0.42 | 0.09 | 0.12 | 0.10 | 0.16 | 0.03 | 0.03 | 0.10 | 0.03 | 0.04 |  |  |  |
| Conditional averaged | $\beta$ | -1.46*** | 0.95* | 0.24* | -0.18 | 0.15 | 0.27 | 0.07 | 0.05 | 0.18 | -0.06 | 0.07 | | | |
|  | SE | 0.17 | 0.42 | 0.09 | 0.10 | 0.11 | 0.27 | 0.10 | 0.10 | 0.21 | 0.11 | 0.11 |  |  |  |

**Appendix 2.** Model selection and model-averaged parameter estimates (β), standard error (SE) for conditional and full averaged model **excluding collared** and legally killed individuals. The models were ranked based on AICc and only the top models (ΔAICc ≤ 2) were kept and used for model averaging. The best model has an AICc of 468.38. The reference in the analyses is “No” for the predictor Collared, “Norway” for Country and “Male” for Sex. *** < 0.001 < ** < 0.01 < * < 0.05

| | | Intercept | Human density | Inbreeding | Gravel road density | Country | Wolf density | Dist. no-breeding zones | Sex | df | $\Delta\text{AICc}$ | weight |
| --- | --- | --- | --- | --- | --- | --- | --- | --- | --- | --- | --- | --- |
|  |  | X | X |  | X |  |  |  |  | 4 | 0 | 0.12 |
|  |  | X | X |  |  |  |  |  |  | 3 | 0.20 | 0.11 |
|  |  | X | X |  | X | X |  |  |  | 5 | 0.33 | 0.10 |
|  |  | X | X | X |  |  |  |  |  | 4 | 0.72 | 0.09 |
|  |  | X | X | X | X |  |  |  |  | 5 | 0.85 | 0.08 |
|  |  | X | X |  |  | X |  |  |  | 4 | 1.06 | 0.07 |
|  |  | X | X | X | X | X |  |  |  | 6 | 1.24 | 0.07 |
|  |  | X | X | X |  | X |  |  |  | 5 | 1.57 | 0.06 |
|  |  | X | X |  |  |  |  | X |  | 4 | 1.60 | 0.06 |
|  |  | X | X |  | X |  |  |  | X | 5 | 1.75 | 0.05 |
|  |  | X | X |  | X |  |  | X |  | 5 | 1.79 | 0.05 |
|  |  | X | X |  |  |  | X |  |  | 4 | 1.86 | 0.05 |
|  |  | X | X |  | X |  | X |  |  | 5 | 1.88 | 0.05 |
|  |  | X | X |  |  |  |  |  | X | 4 | 1.97 | 0.05 |
| Full model averaged | $\beta$ | -1.34*** | 0.23* | -0.04 | 0.10 | 0.11 | $6.10^{-3}$ | $8 \times 10^{-3}$ | 0.01 | | | |
|  | SE | 0.23 | 0.10 | 0.09 | 0.13 | 0.24 | 0.04 | 0.04 | 0.08 |  |  |  |
| Conditional averaged | $\beta$ | -1.34*** | 0.23* | -0.13 | 0.18 | 0.37 | 0.06 | 0.08 | 0.12 | | | |
|  | SE | 0.23 | 0.10 | 0.12 | 0.12 | 0.31 | 0.12 | 0.12 | 0.23 |  |  |  |

**Appendix 3**. Supplementary method for the correction of missing value for snow depth, which was included as environmental factor in our analysis.

From the daily snow depth data extracted from the database SMHI (www.smhi.se) for the weather stations in Sweden and from website seklima [81] using the data from met.no for Norway, some corrections of missing values have been implemented according to the SMHI recommendations, with the following rules. As snow depth measurements are so far always made manually, when there are gaps outside of the winter months (November, March, April, May) the most probable thing is that there had been no observation because there is no snow, so the missing data are replaced by 0. If there is a gap smaller or equal to 14 days, the most probable thing is that it didn’t snow. Thus, if there is an increase (or no difference) in snow depth, the missing values were completed by the latest observed value. If there is a reduction in snow depth, we interpolated linearly and assumed that the snow cover had disappeared or melted at a steady pace. If after these corrections data are still missing for one observation year, the information from this station for this year is discarded.

**Appendix 4.**
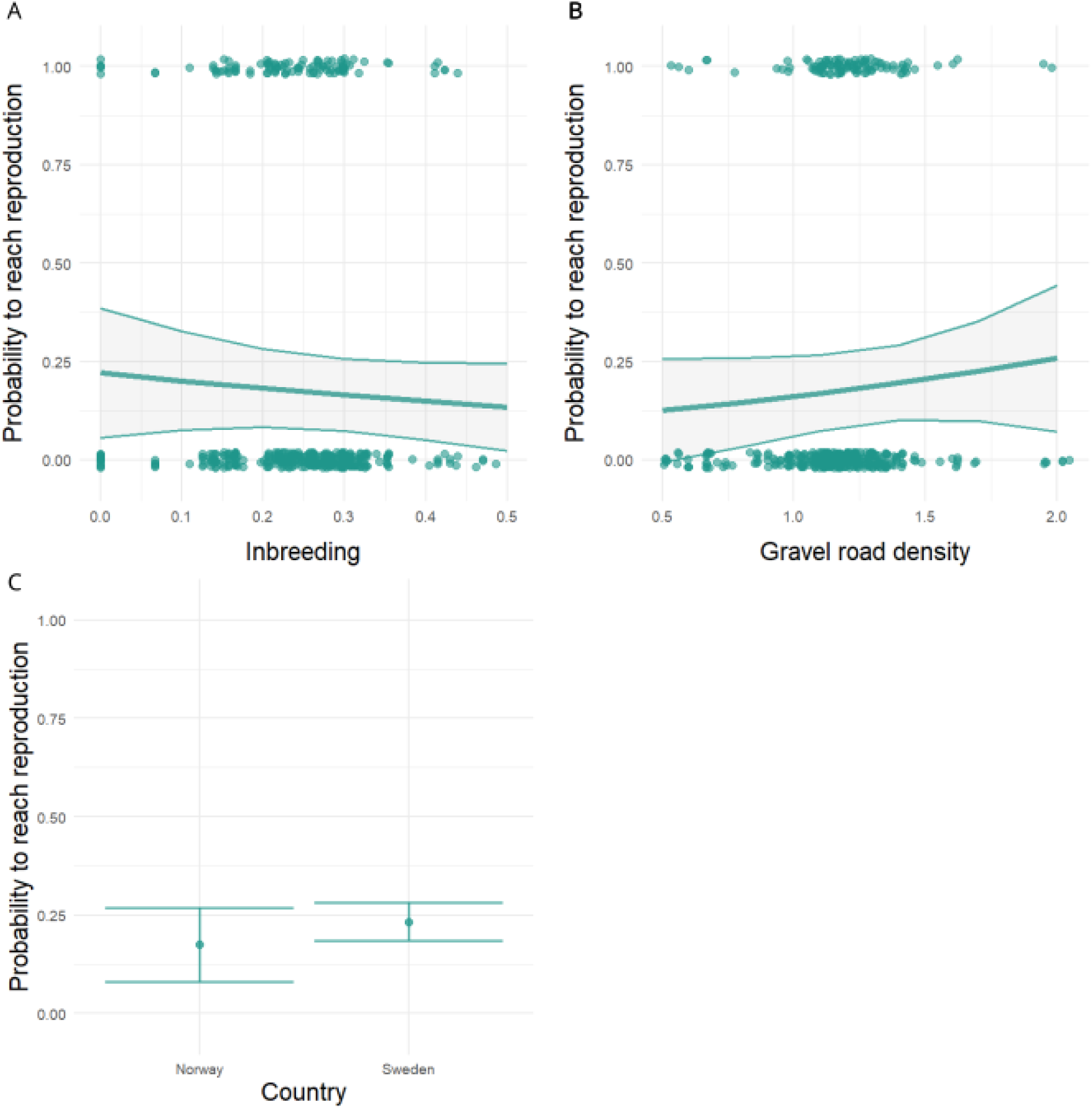
Probability to reach reproduction in relation to A) the inbreeding coefficient, B) the density of gravel road in the natal territory (km) and C) the country of birth. The lines indicate the fitted values, with associated 95% confidence interval from the model-averaged estimates (Table 3). For panels A and B, sex was held constant at “Male”, collared at “No” and birth country at “Norway”, and the other continuous variables coefficients at their mean. The dots in the panel A and B correspond to the observed data points, where their positions on the y-axis represent the actual binary outcomes of reaching reproduction (0 or 1).

